# In vivo optical metabolic imaging of long-chain fatty acid uptake in orthotopic models of triple negative breast cancer

**DOI:** 10.1101/2020.11.02.365288

**Authors:** Megan C. Madonna, Joy E. Duer, Joyce V. Lee, Jeremy Williams, Baris Avsaroglu, Caigang Zhu, Riley Deutsch, Roujia Wang, Brian T. Crouch, Matthew D Hirschey, Andrei Goga, Nirmala Ramanujam

## Abstract

Targeting a tumor’s metabolic dependencies is a clinically actionable therapeutic approach, but identifying subtypes of tumors likely to respond remains difficult. The use of lipids as a nutrient source is of particular importance, especially in breast cancer. Imaging techniques offer the opportunity to quantify nutrient use in preclinical models to aid in the development of new drugs to restrict uptake or utilization of these nutrients. We describe a fast and dynamic approach to image fatty acid uptake in vivo, a tool relevant to study tumor metabolic reprogramming or for studying the effectiveness of drugs targeting lipid metabolism spanning beyond breast cancer and optical imaging alone. Specifically, we developed a quantitative optical approach to spatially and longitudinally map the kinetics of long-chain fatty acid uptake in in vivo murine models of breast cancer using a fluorescently labeled palmitate molecule, Bodipy FL c16. We chose intra-vital microscopy of mammary tumor windows to validate our approach in two orthotopic breast cancer models: a MYC-overexpressing transgenic triple-negative breast cancer (TNBC) model and a murine model of the 4T1 family. Following injection, Bodipy FL c16 fluorescence increased and reached its maximum after approximately 30 minutes, with the signal remaining stable during the 30-80-minute post-injection period. We used the fluorescence at 60 minutes (Bodipy60), the mid-point in the plateau region, as a summary parameter to quantify Bodipy FL c16 fluorescence in subsequent experiments. Using our imaging platform, we observed a two- to four-fold decrease in fatty acid uptake in response to the downregulation of the MYC oncogene consistent with findings from in vitro metabolic assays. In contrast, our imaging studies report an increase in fatty acid uptake with tumor aggressiveness (6NR, 4T07, and 4T1), and uptake was significantly decreased after treatment with a fatty acid transport inhibitor, perphenazine, in both normal mammary pads and in the most aggressive 4T1 tumor model. Our approach fills an important gap between in vitro assays, which provide rich metabolic information but at static time points, and imaging approaches that can visualize metabolism in whole organs, but which suffer from poor resolution.

## Introduction

Cellular metabolism involves a vital network of pathways for homeostasis, growth, and survival. Alterations in these pathways are key features of a variety of diseases and conditions, such as diabetes, obesity, and cancer ^1–3^. In cancer, oncogene activation or loss of tumor suppressors contributes to dysregulated metabolism, causing altered nutrient requirements compared to normal cells ^4,5^. Aerobic glycolysis is frequently upregulated in tumor cells; fluorodeoxyglucose-positron emission tomography (FDG-PET) can measure glucose uptake to stage a patient’s cancer, assess response to therapy, and evaluate progression ^6^. Yet, a steadily growing number of studies suggest that tumors rely on additional energetic sources ^7^, including glutamine, amino acids, and lipids ^8^.

The use of lipids as a nutrient source is of particular importance, especially in breast cancer where an increase in lipid droplet formation is associated with increased tumor aggressiveness ^9,10^. Adipocytes surround invasive breast cancers and serve as professional lipid storing cells, providing a readily available carbon source ^11–14^. Adipocytes surrounding aggressive primary tumors show a loss of lipid content and increased IL-6 expression ^15^, a key inflammatory cytokine previously implicated in migration and invasion of breast cancer cells ^16^. Further, fatty acids are often the preferential carbon source in drug-resistant and residual cancer ^17–19^.

New methods to quantify the use of these energetic sources in preclinical models will aid in the development of new drugs to restrict uptake or utilization of these nutrients. *In vitro* and *ex vivo* assays like the Seahorse Assay and metabolomics quantify multiple metabolic endpoints but at single, static time points and sacrifice spatial information; however, *in vivo* imaging technologies allow for repeated measurements of a single metabolic endpoint for dynamically studying metabolic reprogramming or for studying the effectiveness of potential metabolically targeted therapeutics within the tumor microenvironment with the most common *in vivo* metabolic imaging modality being FDG-PET imaging ^20^.

The ability to directly study fatty acid uptake within the context of an intact tumor-bearing animal is critical for evaluating novel therapeutics that antagonize this pathway or energy source. Previous studies outside of breast cancer have validated the use of an additional radio-labeled PET tracer, 18F-fluoro-6-thia-heptadecanoic acid (18F-FTHA), for *in vivo* fatty acid uptake measurements initially for myocardium imaging ^21^ and more recently in prostate cancer ^22^ and melanoma ^23^, providing more evidence for use of fatty acid uptake as a valuable indicator of study. Though metabolic imaging through PET has proved valuable, especially in the clinic, its millimeter-scale resolution restricts the capturing of a preclinical tumor’s heterogeneity ^24,25^ and limits the ability to test both *in vitro* and *in vivo* models for the development, validation, and translation of metabolically targeted therapies.

Optical imaging allows for the temporal mapping of fatty acid uptake as seen in PET imaging, but at a high resolution for spatial mapping, providing complementary metabolic information to existing techniques. Additionally, optical approaches have the capacity to image several endogenous contrast agents and can be coupled with appropriate exogenous indicators to provide functional and molecular information and allow for high-resolution imaging of both *in vitro* and *in vivo* models of cancer ^26–32^. To quantify exogenous fatty acid uptake, the use of Bodipy FL c16 (4,4-Difluoro-5,7-Dimethyl-4-Bora-3a,4a-Diaza-s-Indacene-3-Hexadecanoic Acid) has been successfully and extensively used *in vitro* to measure fatty acid uptake and, by extension, the cell’s fatty acid (β) oxidation activity ^33^. It is comprised of a fluorophore and a 16 carbon long-chain fatty acid (palmitic acid), the most common saturated fat in animals ^34^. Bodipy FL c16 has also been administered *in vivo* and quantified *ex vivo* by flow cytometry ^35^. The benefit of imaging Bodipy FL c16 is that this would enable both longitudinal and spatial mapping of *in vivo* fatty acid uptake visualization.

Here, we adapted and rigorously validated the fluorescent palmitate analog, Bodipy FL c16, for *in vivo* imaging of fatty acid uptake. This approach can be adapted to optical imaging using a variety of different technologies that differ in terms of depth of penetration, resolution, and field of view. For this study, we specifically developed our methodology using intra-vital microscopy of a mammary window breast tumor model so as to visualize local changes in fatty acid uptake in the precise orthotopic region where the tumor was implanted. We applied our approach to two syngeneic breast cancer models: the MMTV-Tet-O-MYC conditional model of TNBC (MTB-TOM) ^36^ and the transplantable 4T1, 4T07, and 67NR tumors in a mammary carcinoma model ^37^. In the MTB-TOM model, Bodipy FL c16 fluorescence decreased following downregulation of MYC. Conversely, in animals with orthotopic transplantation of the 4T1 family of tumors ^38^, fatty acid uptake increased with tumor metastatic potential. In this same 4T1 family orthotopic model, Bodipy FL c16 fluorescence was significantly decreased with the addition of a small molecule inhibitor, perphenazine, an inhibitor of fatty acid uptake ^39^, exemplifying another use of our approach, the monitoring of metabolic changes following therapy. *In vivo* optical imaging of fatty acid uptake with Bodipy FL c16, connects an important gap in the field of cancer metabolism between *in vitro*-only, single-timepoint assays and *in vivo* PET imaging. Further, these methodologies studies serve as the foundation upon which imaging of fatty acid uptake can be adapted to different types of optical imaging systems.

## Methods

### Ethics Statement

All *in vivo* murine experiments were conducted according to the Duke University Institutional Animal Care and Use Committee (IACUC) (Protocol A072-18-03). All mice were housed in an on-site housing facility with *ad libitum* access to food and water with standard light/dark cycles. Tumor volumes were measured using calipers and calculated as Volume = (Length x Width^2^)/2, where width represents the smallest axis and length the longest axis.

### MYC-overexpressing Murine Model

Viably frozen MTB-TOM tumors, generated from MTB-TOM (MMTV-rtTA/TetO-MYC)^36^ (University of California, San Francisco), were divided into 1-2 mm chunks and transplanted into the 4^th^ right mammary fat pad of 4-week old FVB/N mice (Taconic). Mice were administered 2 mg/ml doxycycline (Research Products International) through their drinking water, to maintain MYC overexpression and tumorigenesis. As tumors reached ~300 mm^3^, mice were imaged, and doxycycline was removed to initiate oncogene down-regulation and tumor regression for 4 days prior to the second imaging session as tumors shrunk by roughly 50% in size. Four FVB/N mice were used for Bodipy imaging while two FVB/N mice were used for TMRE imaging.

### 4T1 Murine Model

The 4T1 and 4T07 cells were acquired from the American Type Culture Collection, and the 67NR cells were generously provided by Dr. Fred Miller (Karmanos Cancer Institute, Detroit, MI) through Dr. Inna Serganova and Dr. Jason Koucher (Memorial Sloan Kettering Cancer Center, New York, NY). Cell lines were passaged every 2 days in RPMI-1640 medium (L-glutamine) with 10% fetal bovine serum (FBS) and 1% antibiotics (Pen Strep). For *in vivo* injection, the abdomen of 6-8 week old female BALB/c (Charles River) was cleaned using 70% ethanol and the 4^th^ right mammary gland was palpated and injected with 100 μL (~50,000 cells) of 4T1 (n = 6 mice), 4T07 (n = 5 mice), or 67NR (n = 5 mice) cells suspended in sterile RPMI-1640 (Corning) with no FBS or antibiotics using a 27G needle. Tumors were imaged at volume of ~150 mm^3^.

### RNA Sequencing

The differential expression data from RNA-sequencing on the MTB-TOM tumors and 4T1 and 67NR cell lines were downloaded from GEO (GSE130922 and GSE104765) ^40,41^. Data is reported as Log2FC after DESeq2 analysis ^42^ in R software.

### Western Blot

Tumor tissue samples were flash-frozen in liquid nitrogen, and protein was extracted via Dounce homogenizer in a cocktail of RIPA buffer (Thermo) and proteinase (Roche) supplemented with phosphatase (Roche) inhibitor. Protein samples were resolved using 4–12% SDS-PAGE gels (Life Technologies) and transferred to nitrocellulose membranes (iBlot 2, Life Technologies). To probe for FATP3 (12943-1-AP, Proteintech, 1:1000) and c-MYC (MYC) (ab32072, Abcam, 1:1000), membranes were first incubated in primary antibody overnight on a 4 °C shaker, then incubated with horseradish peroxidase (HRP)-conjugated secondary antibody. To probe for β-actin (actin), membranes were incubated for one hour at room temperature in HRP-conjugated antibody (sc-47778 HRP, Santa Cruz, 1:10,000). Signals were visualized with ECL (Bio-Rad), and chemiluminescence acquired via Bio-Rad ChemiDoc XRS+. Unsaturated band intensities were quantified using Fiji software.

### Murine mammary window chamber model

Using a previously established procedure, titanium window chambers with 12 mm diameter, No. 2 glass coverslips were surgically implanted over the 4^th^ right mammary gland of 8-12 week old female BALB/c (Charles River) ^43^.

### Imaging probes

For *in vitro* imaging, Bodipy FL c16 (4,4-Difluoro-5,7-Dimethyl-4-Bora-3a,4a-Diaza-s-Indacene-3-Hexadecanoic Acid, Thermofisher) was diluted in sterile RPMI-1640 (Corning) to a final concentration of 1 μM and incubated in each cell plate for 30 minutes. Next, cells were washed and imaged immediately with a standard confocal microscope (Zeiss 780 Upright confocal microscope at the Duke University Light Microscopy Core Facility). A total of 9 representative fields of view across 3 independent plates per cell line were imaged, and each cell line was measured on at least two different days.

For *in vivo* administration, Bodipy FL c16 was diluted to a final concentration of 200 μM in DMSO. This concentration was titrated to achieve a final tissue-level concentration (42.1 nM – 749.9 nM for these studies, calculated from tissue-mimicking phantoms (not shown)) below levels where self-quenching has been observed ^44^ and similar to *in vitro* studies ^33^. TMRE (Tetramethylrhodamine Ethyl Ester, Life Technologies/ThermoFisher) was diluted to a final concentration of 75 μM in sterile PBS according to our previously established protocols ^45^. Both probes were delivered systemically via tail vein injection, and the total volume of each injection was 100 μL.

### Fluorescence microscopy system and metabolic imaging

All mice were fasted for 4 hours (water provided) prior to imaging to ensure a normalized metabolic rate for each animal ^46^. Following fasting, imaging began immediately. The animal was anesthetized using isoflurane (1-1.5% v/v) in room air in an induction chamber. The animal was transferred to a heated (to maintain core body temperature) microscope stage where a background image (laser on) and dark image (laser off) were acquired prior to probe administration to account for signal from sources other than the fluorescence probes. Each animal received 100 μL of 200 μM Bodipy FL c16 in DMSO or 75 μM of TMRE in sterile PBS via tail vein injection and fluorescence images were acquired for 80 minutes. An exposure time of 5 s was used for both Bodipy FL c16 and TMRE images. To account for daily light source variation, all images were background subtracted, beam shape corrected, and calibrated according to a Rhodamine B standard imaged at each imaging session prior to image analysis. Each mouse was imaged only once. The only exception is with our GEM model to track changes in metabolism on dox (MYC-on) compared to no dox (MYC-off) where each mouse was imaged twice, four days apart.

Our previously reported microscope ^45^ was used to capture fluorescence for both of our metabolic endpoints. For Bodipy FL c16 imaging, a 488 nm crystal laser (DL488–100-O, Crystal laser, Reno, NV, USA) was used to excite the probe, followed by a 505 nm longpass dichroic mirror (DMLP505R, Thorlab, USA) to reject the excitation light in the emission channel. Emission signal was collected using a liquid crystal tunable filter (LCTF) (VariSpec VIS-7-35, PerkinElmer, Inc. Waltham, MA, USA, 7 nm bandwidth) programmed to collect at 515 nm and a high-resolution dual-modal charge-coupled device (CCD) (ORCA-Flash4.0, Hamamatsu, Japan). For TMRE imaging, a 555 nm crystal laser (CL555-100-O, Crystal laser, Reno, NV, USA) was used to excite the probe, followed by a 573 nm longpass dichroic mirror (FF573-Di01-25 × 36, Semrock, Rochester, New York, USA) to reject the excitation light in the emission channel. Emission signal was collected using the previously described LCTF (programmed to collect at 585 nm) and CCD. The spectral microscope system was calibrated wavelength by wavelength using a standard lamp source (OL 220 M, S/N: M-1048, Optronic Laboratories, USA).

This system uses a Nikon CFI E Plan Achromat 4x objective (NA = 0.1, Nikon Instruments Inc., USA) for all imaging. This creates a single frame field of view of 2.1 mm x 1.6 mm and a lateral resolution of 2.2 μm, as measured using a 1951 USAF resolution target ^45^. This microscope was controlled by a custom-designed LabVIEW software.

### Fatty acid uptake chemical inhibition

For *in vitro* inhibition, 4T1 cells were treated with 80 μM perphenazine (Sigma Aldrich) dissolved in sterile RPMI-1640 (Corning) for 2 hours prior to *in vitro* staining and imaging according to the methods above. A total of 9 representative fields of view across 3 independent plates per condition were imaged, and each condition was measured on at least two different days.

Separate cohorts of 4T1-bearing (n = 3 mice) or non-tumor (n = 3 mice) animals also received topical treatment with 80 μM perphenazine dissolved in sterile PBS. Following isoflurane anesthesia, the mammary window chamber was removed and 100 μL of perphenazine was applied to the exposed tissue. Following 2 hours of incubation, sterile PBS was used to wash the window, the coverslip was returned to the tissue, and imaging began immediately according to methods above.

### Data processing and statistical analysis

All post-processing and statistical analysis of Bodipy FL c16 fluorescent imaging was performed using MATLAB (MathWorks, USA). Prior to further analysis, each image collected first underwent background and dark noise subtraction by removing the average value of an image collected with the laser source off (dark noise) and removing the average signal imaged prior to fluorescent probe injection (background). Additionally, due to the non-uniformity of illumination of a Gaussian light-source, each image was divided by an image of a uniform phantom to correct for the beam shape. Finally, each image was calibrated using a Rhodamine B standard solution imaged during each imaging session. These methods accounted for autofluorescence, non-uniform illumination, and day-to-day system variations, respectively. The resulting images were used for all statistical comparisons and images displayed in this study. The average intensity values at each imaging timepoint (0-80 minutes) were calculated to generate a time course kinetic curve. Comparisons of these curves were performed using a two-way analysis for variance (ANOVA). Additionally, comparison of average intensities using our summary parameter, Bodipy60, representing the fluorescence that is present 60 minutes post-injection, was performed using a Wilcoxon signed-rank test for paired time points in our MTB-TOM model and using a Wilcoxon rank-sum test followed by a post-hoc Bonferroni correction for multiple comparisons in our 4T1 family of cell lines.

## Results

### In vivo imaging of fluorescently labeled fatty acids in mammary window chambers

Due to their destructive nature, existing technologies that measure long-chain fatty acid uptake within tumor tissues are often limited in their ability to temporally study a single tumor ^47,48^. Fluorescence imaging provides an opportunity to non-invasively study the dynamics of fatty acid metabolism over time, moving past the limitations of a static snapshot of fatty acid levels. To capture longitudinal fatty acid uptake and model a patient treatment cascade, we designed an imaging strategy using Bodipy FL c16. Bodipy FL c16, a commercially available fluorescent palmitate analog, consists of a 16-carbon long-chain fatty acid molecule linked to a Bodipy dye (**Fig. 1A**). Bodipy FL c16 can be delivered systemically via tail vein injection (**Fig. 1B**). Importantly, Bodipy FL c16 only enters cells via long-chain fatty acid transport proteins like normal fatty acids and not through simple cell membrane diffusion ^33^, making Bodipy FL c16 fluorescence a reliable indicator of true fatty acid transport. Once the probe is taken up into the target cell, it accumulates in the cytoplasm, providing an indicator of how rapidly cells are importing fatty acids. Because the 16^th^ carbon of the palmitate is fluorescently labeled, this prevents immediate degradation of the Bodipy dye during fatty acid oxidation (β-oxidation).

**Figure 1:**
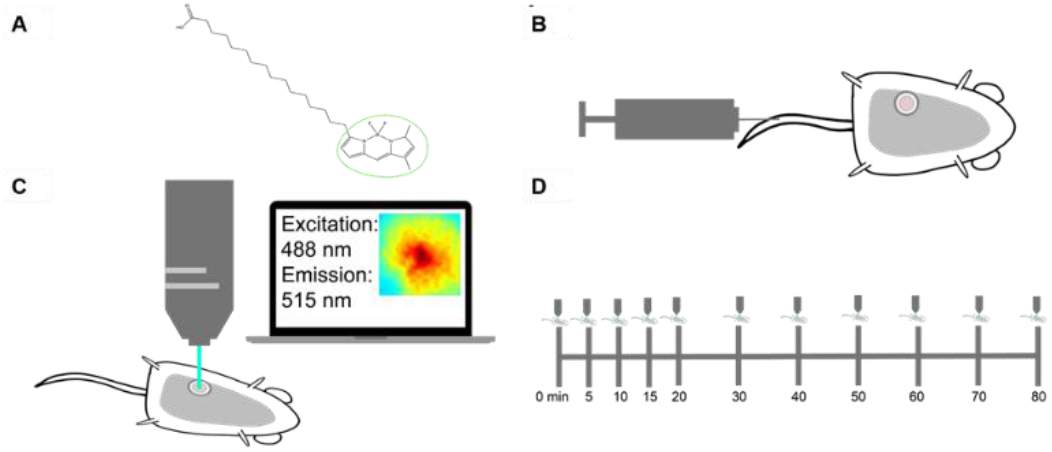
Methods for imaging palmitate uptake through Bodipy FL c16 in mammary window chambers using intravital fluorescence microscopy. **A**, Schematic of fluorescently-labeled palmitate, a 16 carbon-chain fatty acid (Bodipy FL c16). The fluorescent label is marked with a green circle. **B**, Prior to imaging, a titanium mammary window chamber is implanted over the 4^th^ right mammary fat pad of the mouse. 100 μL of 200 μM Bodipy FL c16 is administered via the tail vein for each imaging session. **C**, Fluorescent probe accumulation into the mammary window chamber is measured by exciting the probe at 488 nm and collecting the emitted signal at 515 nm. **D**, Timeline indicating the longitudinal imaging time course for each imaging session.

For intravital imaging, a 12 mm diameter titanium window was implanted over the tumor in the 4^th^ mammary gland of each female mouse. The field of view within the window was excited at 488 nm +/− 5 nm and collected at 515 nm +/− 3.5 nm emission for fluorescence imaging (**Fig. 1C**). To account for endogenous fluorescence differences between tumors, a background and dark image were collected and subtracted from each post-injection image to ensure all reported signal was only due to Bodipy FL c16. Each mouse was imaged sequentially for a total of 80 minutes post-Bodipy FL c16 injection **(Fig. 1D).**

### Bodipy FL c16 uptake differentiates MYC oncogene signaling in tumors

Previous studies found tumors that overexpress the MYC oncogene, such as receptor triple-negative breast cancers (TNBC), have increased uptake of stable isotope-labeled fatty acids in a MTB-TOM model ^49^. We chose to assess our optical imaging technology using the same model, where MYC expression is activated by doxycycline (dox) ^36,49^; modulation of MYC through dox allows for temporal assessment of metabolism as tumors grow while on dox and regress after dox removal. To image Bodipy FL c16 uptake during murine tumor growth and regression, 100 μL of 200 μM Bodipy FL c16 was injected via the tail vein and fluorescence microscopy images (field of view: 2.1 mm x 1.6 mm) of the mammary tumor were captured over a span of 80 minutes post-injection while mice were receiving ad libitum dox (tumor growth with MYC-on) or four days after withdrawal of dox (early tumor regression with MYC-off ^50^). **Fig. 2A** shows representative pre- and post-injection images from growing and regressing tumors. Following MYC inhibition for 4 days, tumors regressed in size by ~2-fold (320 vs. 173 mm^3^; p<0.05), and Bodipy FL c16 uptake is significantly diminished, ~ 3-fold less at peak (**Fig. 2B;** p < 0.05) compared to MYC-on tumors. Normalizing each kinetic curve to its respective maximum fluorescence (**Fig. 2C**) shows that both the MYC-on and MYC-off groups accumulate Bodipy FL c16 for approximately 30 minutes, with signal remaining stable during the 30-80-minute post-injection period. We used the fluorescence at 60 minutes (Bodipy60), the mid-point in the plateau region, as a summary parameter to quantify Bodipy FL c16 fluorescence in subsequent experiments. Thus, MYC-dependent uptake of long-chain fatty acids can be visualized with Bodipy FL c16 in accordance with previously reported stable isotope uptake studies in this model ^49^.

**Figure 2:**
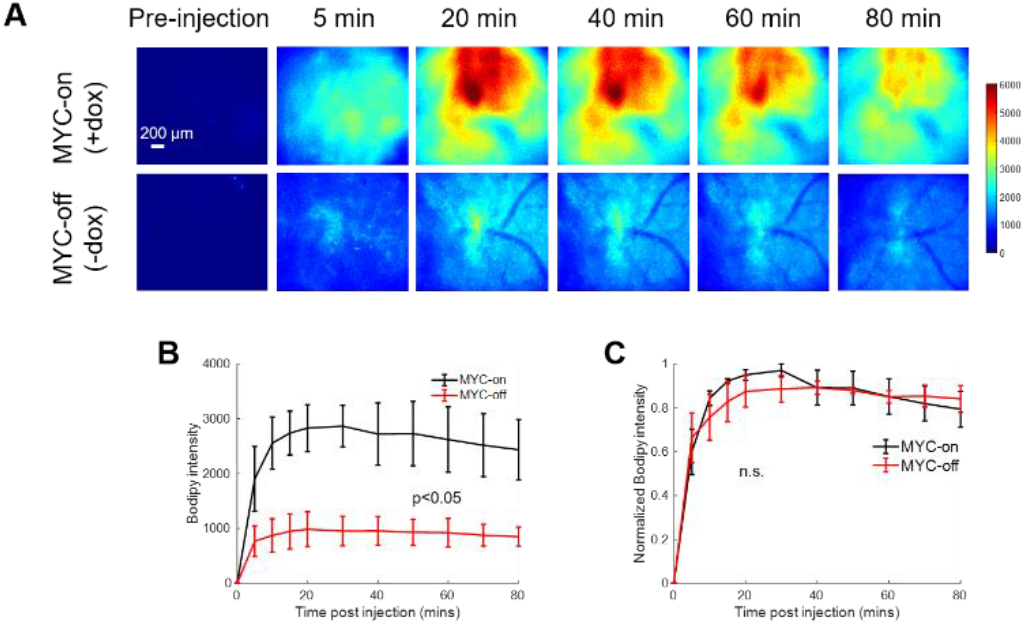
Bodipy FL c16 fluorescence peaks and then plateaus 30 minutes post-injection and MYC downregulation (MYC-off) results in a significant decrease in Bodipy FL c16 uptake. Bodipy FL c16 uptake was measured in mammary tumors overexpressing the oncogene MYC (+ dox) and mammary tumors in which MYC was downregulated (-dox). **A**, Representative images of Bodipy FL c16 fluorescence images over 80 minutes in a mammary tumor with MYC-on (+dox) and MYC-off (four days after dox was removed). Baseline images were acquired prior to Bodipy FL c16 injection and its average was subtracted from the images acquired after Bodipy FL c16 administration. Scale bar = 200 μm. **B,** Average Bodipy FL c16 kinetic curves for the MYC-on (+dox) and MYC-off (-dox) groups. **C,** For a given mouse, the kinetic curve was normalized to its peak Bodipy FL c16 fluorescence and normalized kinetics were then averaged for each group. Error bars represent standard error. (Sample size: MYC-on = 4 mice, MYC-off = 4 mice). The mid-point of the plateau, 60 minutes, was used to represent the Bodipy FL c16 (Bodipy60) fluorescence for subsequent analyses. Statistical differences in mean kinetic curves were determined using a two-way Analysis of Variance (ANOVA) test.

### MYC overexpression corresponds to increased fatty acid transport protein expression

Long-chain fatty acid (LCFA) uptake and trafficking within cells can be mediated by multiple receptors and carriers. Therefore, the increased Bodipy FL c16 uptake observed in MYC-on tumors may be mediated by multiple effector molecules. To evaluate which effector molecules are important for fatty acid uptake in MYC-driven tumors, we characterized the effect of MYC expression on cell surface transport proteins which have been reported to mediate LCFA uptake (**Supplemental Figure 1A**). CD36 and the SLC27 gene family responsible for creating the fatty acid transport proteins (FATPs) have been shown to be critical in both the uptake and activation of long-chain fatty acids for fatty acid oxidation (β-oxidation) ^51,52^. We analyzed RNA sequencing data of MTB-TOM tumors ^40^ to investigate how regulating MYC expression affects CD36 and FATP-family expression. When MYC was turned off through dox withdrawal, at an RNA level, *Slc27a3* was the mRNA sequence in the family that decreased in expression, though not significantly (**Supplemental Figure 1A**). However, MYC may indirectly regulate protein abundance by altering ribosome biogenesis and translation pathways ^53–55^. The mRNA *Slc27a3* sequence specifically codes for FATP3; thus, we examined FATP3 protein expression in multiple MTB-TOM tumors. We found concomitant decreases in MYC and FATP3 protein expression when dox is removed during early tumor regression (**Fig. 3A**; full blot **Supplemental Figure 1B**); FATP3 protein expression was diminished by ~ 75% with the removal of dox (MYC-off) (**Fig. 3B**; p<0.05). While it is possible that multiple mechanisms contribute to increased Bodipy FL c16 uptake in MYC-on tumors, which are beyond the scope of the present study, we have identified at least one FATP-family member, FATP3, whose expression is correlated to MYC in a model of TNBC.

**Figure 3:**
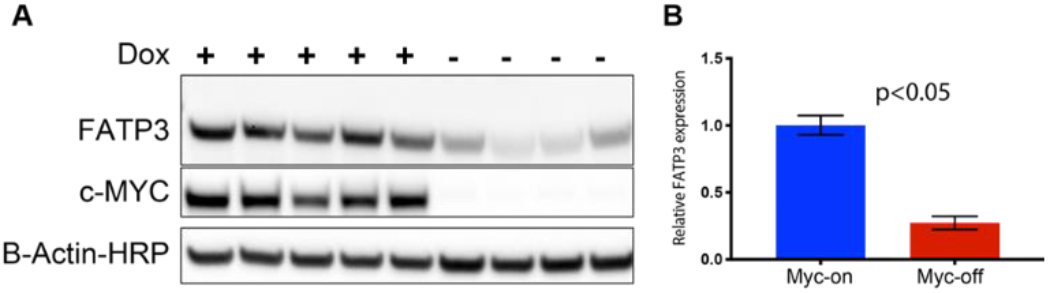
Loss of MYC expression results in decreased FATP3. Fatty acid transport protein 3 (FATP3) expression is related to MYC expression. **A,** Western blots of FATP3 and MYC expression in tumors with (+) or without (-) dox for 4 days confirm FATP3 decreases as MYC expression decreases. Cropped western blots are shown; full blots can be found in Supplemental Figure 1. **B,** Significant differences observed in FATP3 expression between MYC-on (+ dox) vs. MYC-off (-dox) tumors. (Sample size: + dox = 5 mice, - dox = 4 mice). Statistical difference in expression was performed with a Student’s t-test.

### Longitudinal imaging reveals modulation of MYC expression changes mitochondrial metabolism

With the importance of fatty acids in these MYC-on tumors established, we next explored how our method can longitudinally track metabolic changes in each animal following MYC downregulation. To test this, we imaged animals while MYC-on (+dox) tumors and 4 days later, injected more dye and imaged again while MYC-off (-dox for 4 days) tumors (**Fig. 4A**). We saw a significant decrease in average Bodipy60 fluorescence for MYC-off tumors compared to MYC-on tumors (p<0.05) using a Wilcoxon signed-rank test (**Fig. 4B)**. Notably, we observed inter-tumoral heterogeneity in lipid accumulation but nevertheless saw each mouse’s individual decrease in Bodipy60 following MYC-downregulation (dox withdrawal). Additionally, due to the nature of the imaging approach, spatial information across the window and intensity distributions can be captured through probability density functions (PDFs) across each experimental group (**Fig. 4C**). PDFs represent the pixel distribution of all Bodipy FL c16 fluorescence for each imaging group. The PDF of MYC-off (-dox) is significantly left-shifted relative to the MYC-on (-dox) group by Kolmogorov–Smirnov (KS) testing (p<0.05), signifying a decrease in fluorescence (fatty acid uptake) in the tissue.

**Figure 4:**
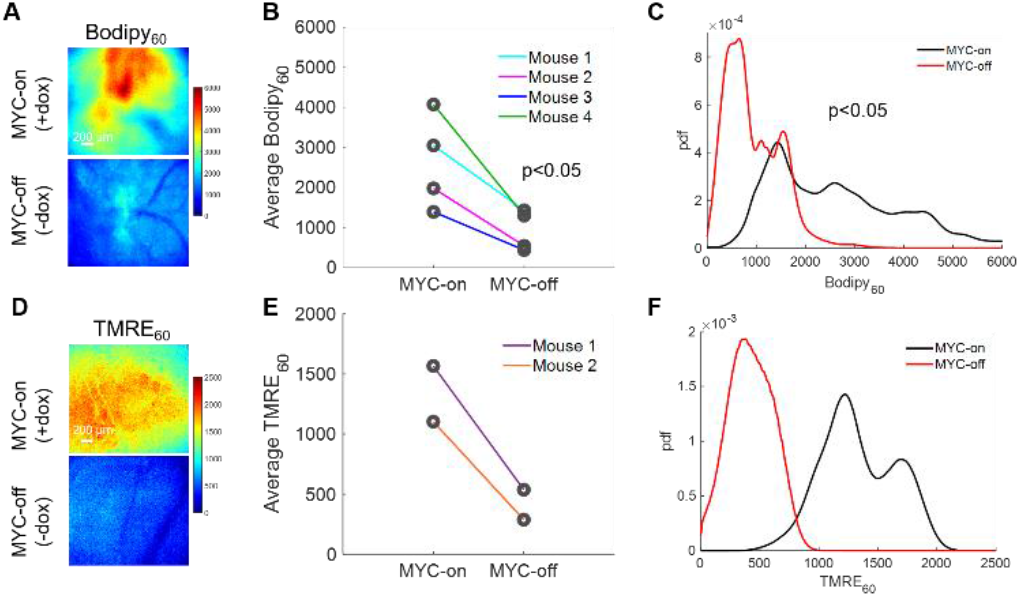
Intravital imaging demonstrates MYC dependence on fatty acid uptake and mitochondrial oxidation. **A**, Representative Bodipy_60_ images for MYC-on (+dox) and MYC-off (-dox) tumors. **B**, Average Bodipy_60_ fluorescence on day 0 (MYC-on) and day 4 (MYC-off) for four tumor-bearing mice. **C**, Bodipy_60_ Probability density functions for MYC-on (+dox) and MYC-off (-dox) across all pixels and all mice in each group. **D**, Representative TMRE_60_ images for MYC-on (+dox) and MYC-off (-dox). **E**, Average TMRE_60_ fluorescence on day 0 (MYC-on) and day 4 (MYC-off) for two tumor-bearing mice. **F**, TMRE_60_ PDFs for MYC-on (+dox) and MYC-off (-dox) across all pixels and all mice in each group. Scale bar = 200 μm. Statistical differences in average fluorescence between groups were determined using a Wilcoxon signed-rank test, and statistical differences in fluorescent pixel distributions were determined using a KS test.

As LCFAs serve as an important carbon source for oxidative phosphorylation, after being broken down through β-oxidation, in MYC-on tumors ^49,56^, understanding how decreased availability of fatty acids through the reduction in MY C expression may affect oxidative phosphorylation would permit for dynamic *in vivo* assessment of tumor overall metabolic reprogramming. Previously, our group has established a method to optically measure oxidative phosphorylation *in vivo* in preclinical models via tetramethylrhodamine ethyl ester perchlorate (TMRE) ^32^. This cationic molecule accumulates in proportion to the mitochondrial membrane potential ^57^, which serves as a surrogate for oxidative phosphorylation rates (i.e., increased mitochondrial membrane potential correlates with increased levels of oxidative phosphorylation). Due to the unknown optical, biological, or chemical crosstalk between Bodipy FL c16 and TMRE probes, a separate cohort of mice was used to investigate TMRE uptake in MYC-on and MYC-off tumors. Specifically, 100 μl of 75 μM TMRE was injected via the tail vein and fluorescence images were captured from the mammary window over a span of 60 minutes according to our previously validated methods ^32^. We observed TMRE60 (TMRE fluorescence 60-minutes post-injection) images for MYC-on and MYC-off breast tumors (**Fig. 4D)**. Roughly a 3-fold decrease in TMRE60 fluorescence was observed following MYC downregulation (**Fig. 4E)**. Additionally, the distribution of TMRE60 fluorescence shifted left in MYC-off (-dox) tumors relative to MYC-on (+dox) tumors, indicating a decrease in oxidative phosphorylation (**Fig. 4F**). Here, we demonstrate the dynamic assessment of both LCFA uptake and oxidative phosphorylation in MYC-driven breast tumor models during tumor growth and early regression.

### Fatty acid uptake increases with metastatic potential

Because alterations in tumor metabolism can influence how aggressive a tumor is ^9,10^, we applied our Bodipy FL c16 imaging technique to test whether our method can distinguish more and less aggressive tumors. The 4T1 tumor family, consisting of 67NR, 4T07, and 4T1 cell lines that arose from the same TNBC parental tumor ^38^, is known for the differing metastatic potential of each cell line ^37^. The 4T1 tumor cells are considered the most aggressive because they have the highest rate of metastases, with secondary nodules often forming in the lungs and liver; the 4T07 tumor cells can mobilize from the primary tumor site but are only found in the blood, lymph nodes, and sometimes the lung, where they often fail to proliferate; the 67NR tumor cells are not detected in the blood, lymph nodes, or lungs ^37,38^. Previous work has demonstrated 4T1 and 4T07 cells have increased oxidative phosphorylation compared to 67NR cells as well as the 4T1 cells’ increased glycolytic flux compared 67NR cells observed in an *in vitro* setting ^58^.

Both *in vitro* studies on cell monolayers and *in vivo* studies on mammary tumors in a window chamber model were performed here. Briefly, cell monolayers were incubated with 1 μM Bodipy FL c16 for 30 minutes prior to being washed and imaged immediately with a confocal microscope. Representative images are shown in **Fig. 5A** and differences in fluorescence quantified in **Fig. 5B.** Bodipy FL c16 uptake in 4T1 and 4T07 cell lines are significantly higher than that in the 67NR cell line (p<0.05). These same trends are observed *in vivo* as shown in **Fig. 5C-D.** The 4T1, 4T07, and 67NR tumors were all grown to 5 mm x 5 mm in the mammary fat pad of female BALB/c mice, which maintain a fully intact immune system. Representative Bodipy60 images for each group are shown in **Fig. 5C**. The average Bodipy60 for each animal increased in 4T1 and 4T07-bearing mammary windows compared to 67NR-bearing mammary windows (p<0.05) (**Fig. 5D).** Furthermore, we probed a recently published RNA-sequencing dataset and found 4T1 cells have higher *Myc* mRNA when compared with 67NR cells ^41^. While *Slc27a3* mRNA expression followed the trend of *Myc* expression, it was not significantly different between 67NR and 4T1 cells (**Supplemental Figure 2**). These data show that the more metastatic 4T1 and 4T07 primary tumors have increased fatty acid uptake compared to non-metastatic tumors.

**Figure 5:**
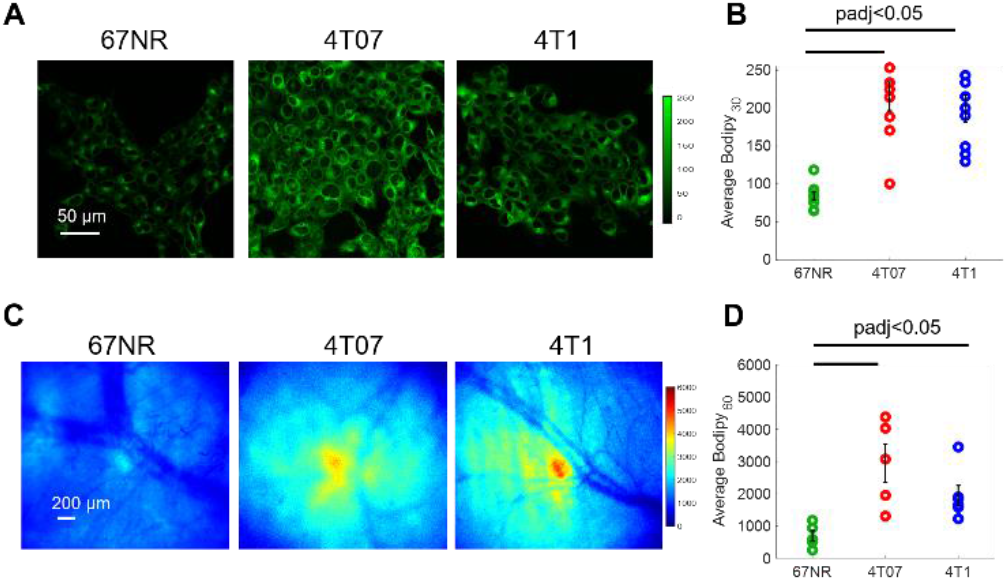
Bodipy_60_ is increased in metastatic-prone tumors Bodipy_60_ fluorescence of sibling breast tumor lines (4T1, 4T07, and 67NR) *in vitro* and *in vivo.* **A**, Representative Bodipy FL c16 images of cells stained with Bodipy FL c16 for 30 minutes for each cell type. Scale bar = 50 μm. **B**, Average Bodipy FL c16 fluorescence per cell line. (Sample size: all groups = 9 fields of view across 3 plates). **C**, Representative Bodipy_60_ images of 67NR, 4T07, and 4T1 tumors *in vivo.* Scale bar = 200 μm. **D**, Average Bodipy_60_ fluorescence for each tumor line. (Sample size: 4T1 = 6 mice, 4T07 = 5 mice, 67NR = 5 mice). Error bars = standard error. Statistical differences in average fluorescence between groups were determined using a Wilcoxon rank-sum test followed by a post hoc Bonferroni correction for multiple comparisons.

### Using Bodipy FL c16 to determine drug efficacy

Finally, we demonstrated that Bodipy FL c16 imaging can be applied to study *in vivo* efficacy of drugs that attenuate LCFA uptake. We tested whether perphenazine, a drug previously shown to inhibit LCFA uptake ^39,59^, can reduce fatty acid uptake in 4T1 tumors. Perphenazine, an FDA approved drug used to treat patients with schizophrenia, binds to a subset of FATPs and reduces LCFA uptake *in vitro* and in a yeast-based functional screen^39,59^.

We first imaged Bodipy FL c16 uptake *in vitro* using a confocal microscope following a 2-hour incubation with 80 μM perphenazine (**Fig. 6A**) per standard protocols^39^. Bodipy FL c16 uptake was significantly decreased in 4T1 cells treated with perphenazine (**Fig. 6B**) (p<0.05). To monitor fatty acid uptake following perphenazine treatment *in vivo*, mammary window chambers were established, as previously described, above 4T1 tumors. On the day of imaging, the mammary window coverslip was removed and 80 μM perphenazine was topically applied to the exposed tissue for 2 hours ^39^. Following treatment, the window was rinsed with sterile PBS, followed by our standard Bodipy FL c16 imaging protocol. Representative Bodipy_60_ images of a 4T1 tumor with and without perphenazine treatment are shown in **Fig. 6C.** Short-term treatment with perphenazine significantly decreased Bodipy_60_ in 4T1 tumors (**Fig. 6D**) (p<0.05). We observed comparable results with *in vivo* non-tumorous mammary tissue treated with and without perphenazine (**Supplementary Figure 3**).

**Figure 6:**
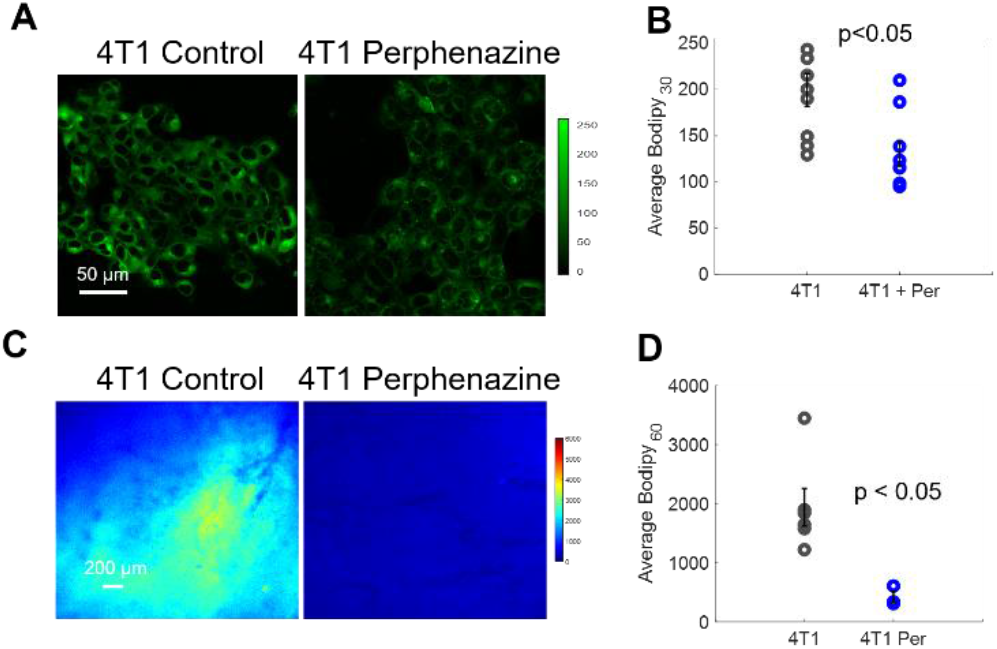
Bodipy_60_ decreases with the inhibition of fatty acid uptake. **A**, Representative Bodipy Fl c16 images of 4T1 cells treated with PBS (4T1 control) or 80 μM of perphenazine (4T1 Perphenazine) for 2 hours prior to staining with Bodipy Fl c16 for 30 minutes. Scale bar = 50 μm. **B**, Average Bodipy FL c16 fluorescence per experimental group. (Sample size: all groups = 9 fields of view across 3 plates). **C**, Representative Bodipy_60_ images of 4T1 tumors. Scale bar = 200 μm **D**, Average Bodipy_60_ fluorescence for each experimental group. (Sample size: Control = 6 mice, Perphenazine = 3 mice). Per = perphenazine treated. Error bars = standard error. Statistical differences in average fluorescence between groups were determined using a Wilcoxon rank-sum test.

## Discussion

Our and others’ prior work indicate evidence of fatty acid dependencies in certain cancers ^22,49,60–64^. However, there are surprisingly few technologies that can directly compare tumor lipid uptake *in vivo*, especially at length scales that can fill the gap between *in vitro* assays and whole-organ imaging. In the work presented here, we addressed this shortcoming in the literature through the development and validation of a fluorescence-based, longitudinal fatty acid uptake imaging approach in two distinct *in vivo* murine models. With this approach, we found increased fatty acid uptake in MYC-high tumors and tumors with high metastatic potential. Additionally, we demonstrate fatty acid inhibition with a reported decrease in fluorescent signal following the addition of perphenazine, a known fatty acid inhibitor. Taken together this approach allows for spatial and temporal mapping of cancer cell fatty acid uptake in local tumor microenvironments.

We tested our imaging approach in a MYC model, which has previously been shown to take up fatty acids from the circulation, validating the ability of our method to quantify fatty acid uptake in a regulated system. Bodipy FL c16 imaging, however, has the added advantage that it permits dynamic temporal imaging as MYC-driven tumors form and regress. The MTB-TOM model showcases two critically needed features for clinically relevant studies of fatty acid uptake: 1) longitudinal metabolite tracking in a single animal for intra-animal decreases in fatty acid uptake following oncogene downregulation; and 2) providing a link between fatty acid uptake and tumor aggressiveness. This decreased uptake also correlated with a visible reduction in tumor size ^50^.

Longitudinal imaging of Bodipy FL c16 led to the finding that MYC-dependent tumors have decreased fatty acid uptake following MYC downregulation, which serves as a proof of concept for future testing of MYC specific therapeutic interventions. We link FATP3 protein expression with MYC expression. While our data suggest that treatment with inhibitors of FATP3 or knocking down this LCFA carrier protein would drive anti-tumor effects under the setting of MYC-high tumors, we did not directly test it here, as it was beyond the scope of the present study. Future studies can be carried out to test the role of FATP3 in breast cancer.

Previous studies have implicated increased both fatty acid uptake and β Oxidation in overall tumor aggressiveness and metastatic potential beyond MYC-inducible models ^15,65–69^. The tumors of the 4T1 family of cell lines, which vary in metastatic potential (4T1 > 4T07 > 67NR), were selected to assess whether our technique could associate differences in fatty acid uptake across tumors of varying metastatic potential ^38^. Not only was Bodipy FL c16 fluorescence indicative of metastatic potential (higher in 4T1 and 4T07 vs. 67NR), but also, through an existing dataset, we show *Myc* mRNA expression is higher in 4T1 cells in comparison with 67NR cells. This is consistent with previous findings that breast cancer patients with the highest *Myc* gene signature are most likely to have disease recurrence ^70^. Aside from MYC-driven tumors alone, fatty acid oxidation in TNBC and its respective metastasis have recently been linked to the activation of the SRC oncoprotein ^69^. SRC expression specifically has been shown to be elevated in both 4T1 and 4T07 tumors compared to 67NR tumors and the knockdown of *Src* significantly inhibited the metastatic burden of 4T1 tumors ^71,72^. The increased fatty acid uptake in all aggressive tumor lines suggests a metabolic switch associated with the use of exogenous fatty acids in the spread of metastatic disease. Future studies can genetically manipulate the 4T1 cells to reduce their metastatic potential and examine whether lipid uptake could be used as a complementary readout for reduced aggressiveness. This idea is supported by previous studies implicating fatty acid uptake and oxidation as key to promote tumor cell migration ^67,73,74^.

Intravital Bodipy FL c16 imaging can also report on the activity of fatty-acid-uptake-targeted therapies, such as perphenazine, both *in vivo* and *in vitro*. While we do not know the exact mechanism, the drug likely targets the FATP family. FATP (1-6) have roles in fatty acid uptake and/or activation in cells, so our group sought to image its effects on Bodipy FL c16’s uptake kinetics ^59^. Prior research in yeast demonstrated perphenazine’s ability to block FATP2 ^75^; however, there is likley some inhibitory effect on other FATP receptors. Our data supports additional studies on perphenazine in the future, where our method could be used to examine drug-dosing and minimal doses to target fatty acid uptake and oxidation in cancer, in order to reduce side effects. Though only one drug is tested here, the platform can be applied to a wide variety of fatty acid uptake or β oxidation inhibitors through either topical or systemic administration.

This work demonstrates that optical imaging of Bodipy FL c16 imaging is important for lipid metabolism research. These studies are paramount to the successful translation of targeted metabolic therapies for TNBC and the plethora of other tumors that are dependent on fatty acids ^61,69,76^. The drug screening protocol demonstrated here could also be applied to image the liver or brain through abdominal or cranial window chamber models, respectively ^77^, to capture diseased livers ^78^ or brain metastases ^79^, which have reportedly shown a dysregulated lipid metabolism. Additionally, the methodology developed here can be extended to other optical imaging modalities such as IVIS or fluorescence spectroscopy systems for non-invasive monitoring of solid tumors. Collectively, this technique can provide a quantitative measurement for fatty acid uptake across various disease and imaging models to screen potential therapeutics.

We provide evidence supporting the use of fatty acid uptake with other validated metabolic imaging techniques (TMRE) to indicate effects on fatty acid oxidation in these tumors through the addition of mitochondrial membrane potential imaging ^32^. These results are consistent with our prior work ^49^. MYC overexpression can reprogram tumor metabolism in multiple ways, including uptake and utilization of multiple metabolites that can contribute to increased oxidative phosphorylation including LCFA, glucose, and glutamine ^80^. Through *in vivo* optical imaging, metabolic phenotyping of MYC driven cancers is feasible. In addition, fluorescence imaging can capture multiple endpoints through spectral unmixing as we have shown previously, where membrane potential and glucose uptake can be reported in the same imaging session ^45^. This is beneficial compared to PET imaging where dual-tracer PET imaging studies typically require a waiting period of at least 24 hours in between imaging of each probe due to the similar nature of PET signal regardless of the isotope-labeled biomolecule^81–83^.

As the growing literature implicates fat as a key carbon source to many cancers, there is a need for a validated method to non-invasively image fatty acid uptake *in vivo.* In breast cancer alone, for example, exogenous fatty acid uptake and oxidation have been shown to be increased in primary TNBC ^80^ and following therapy resistance in Her2 breast cancer ^18^ as well as in residual disease across breast cancer subtypes ^19^. Pinpointing particular cancers with alterations in lipid metabolism may provide clues to targeting oncogenic metabolism in the clinic. We present a method for *in vivo* optical imaging of fatty acid uptake to metabolically phenotype *in vivo* tumors within their microenvironment. This technique is complementary to existing *in vitro* metabolic assays and *in vivo* PET imaging and can be used to detect and inform on alterations of tumor metabolism in preclinical studies and screen and validate potential therapy efficacy for translation to future clinical studies.

## Acknowledgements

Thank you to Dr. Fred Miller; Karmanos Cancer Institute, Detroit, MI for the original identification and dissemination of the 67NR cell line. Many thanks to Dr. Jason Koutcher, Dr. Inna Serganova, and Dr. Natalia Kruchevsky at Memorial Sloan Kettering Cancer Center for generously providing and arranging delivery of the 67NR cell line to our lab. Thank you to Amy Frees Martinez for assistance in statistical analysis. Thank you to Roman Camarda for training and assistance in animal surgeries. We would also like to acknowledge Ken Young for protocol assistance. Finally, many thanks to Jihong Lee for assistance in preliminary experiments.

This work is supported by the Duke Medical Imaging Training Program NIH Grant T32-EB001040, 1F31CA243194, R01EB028148-01, F32CA243548, 1F31CA243468, R01CA223817 and CDMRP W81XWH-18-1-0713 and W81XWH-16-1-0603. The funders had no role in study design, data collection and analysis, decision to publish, or preparation of the manuscript.

