## Supplemental Data for "In vivo optical metabolic imaging of long-chain fatty acid uptake in orthotopic models of triple negative breast cancer"

---

### Supplemental Data

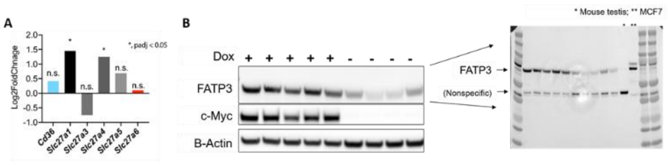


**Supplemental Figure 1: RNA sequencing and full western blot confirm removal of dox is correlated with decreased *SLC27a3*/FATP3 expression. A**, RNA sequencing data from MTB-TOM tumors comparing 4 days after dox withdrawal (regression) with MYC-on tumor. (Sample size: n = 3). **B**, Western blots of FATP3 and MYC expression in tumors with (+) or without (-) dox for 4 days confirm FATP3 decreases as MYC expression decreases. (Sample size: + dox = 5 mice, - dox = 4 mice). Uncropped blots are shown to the right, with mouse testis (*) and breast cancer line MCF7 (**) as expression controls.

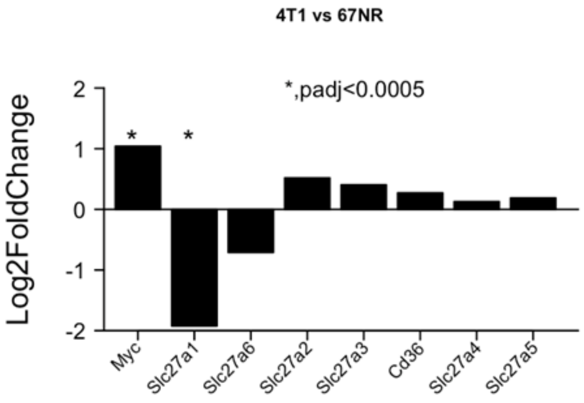


**Supplemental Figure 2:** RNA sequencing data analysis of murine breast cancer models (4T1 - highly metastatic and 67NR - poorly metastatic) shows differential regulation of MYC. Here, an increased fold change corresponds with a gene expressed more in 4T1 cells than 67NR cells. (Sample size: n = 3).

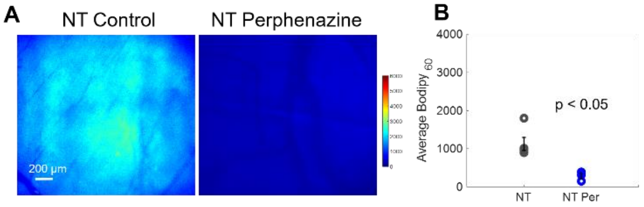


**Supplemental Figure 3: Bodipy$_{60}$ decreases with the inhibition of fatty acid uptake in normal, mammary gland tissue A**, Representative Bodipy$_{60}$ images non-tumor (NT). Scale bar = 200 µm. **B**, Average Bodipy$_{60}$ fluorescence for each experimental group. (Sample size: Control = 5 mice, Perphenazine = 3 mice). Error bars = standard error. Statistical differences in average fluorescence between groups were determined using a Wilcoxon rank-sum test.
